# LFMM 2.0: Latent factor models for confounder adjustment in genome and epigenome-wide association studies

**DOI:** 10.1101/255893

**Authors:** Kevin Caye, Basile Jumentier, Olivier François

## Abstract

**Motivation:** Genome-wide, epigenome-wide and gene-environment association studies are plagued with the problems of confounding and causality. Although those problems have received considerable attention in each application field, no consensus have emerged on which approaches are the most appropriate to solve this problem. Current methods use approximate heuristics for estimating confounders, and often ignore correlation between confounders and primary variables, resulting in suboptimal power and precision.

**Results:** In this study, we developed a least-squares estimation theory of confounder estimation using latent factor models, providing a unique framework for several categories of genomic data. Based on statistical learning methods, the proposed algorithms are fast and efficient, and can be proven to provide optimal solutions mathematically. In simulations, the algorithms outperformed commonly used methods based on principal components and surrogate variable analysis. In analysis of methylation profiles and genotypic data, they provided new insights on the molecular basis of diseases and adaptation of humans to their environment.

**Availability and implementation:** Software is available in the R package lfmm at https://bcm-uga.github.io/lfmm/.

## 1 Introduction

Association studies have been extensively used to identify candidate genes or molecular markers associated with disease states, exposure levels or phenotypic traits. Given a large number of target variables, the objective of those studies is to test whether any of the variables exhibits significant correlation with a primary variable of interest. The most common association studies are genome-wide association studies (GWAS) that focus on single-nucleotide polymorphisms (SNPs) by examining genetic variants in different individuals [2]. In recent years, other categories of association studies have emerged and become important. Of specific interest, epigenome-wide association studies (EWAS) measure DNA methylation levels in different individuals to derive associations between epigenetic variation and exposure levels or phenotypes [35]. Gene-environment association studies (GEAS) test for correlation between genetic loci and ecological variables in order to detect signatures of environmental adaptation [36].

Although they could bring useful information on the causes of diseases or on biological functions, association studies suffer from the problem of confounding. This problem arises when there exist unobserved variables that correlate both with primary variables and genomic data [42]. Confounding inflates test statistics, and early approaches consisted of introducing inflation factors to correct for the bias [5]. Statistically, inflation factors represent an empirical null-hypothesis testing approach, which is frequently used in gene expression studies [10]. GWAS have addressed the confounding issue by including known or inferred confounding factors as covariates in regression models. A prominent GWAS correction method is principal component analysis (PCA) which adjusts for confounding by using the largest PCs of the genotypic data [32]. A drawback of the approach is that the largest PCs may also correlate with the primary variables, and removing their effects can result in loss of statistical power. In gene expression studies where batch effects are source of unwanted variation, alternative approaches to the confounder problem have been proposed. These methods are based on latent factor regression models, also termed surrogate variable analysis (SVA) [26, 7]. Latent factor models have also been considered in GEAS (LFMM, [13]) and in EWAS for dealing with cell-type composition without reference samples (RefFreeEWAS, [21, 39]). Latent factor models employ deconvolution methods in which unobserved batch effects, ancestry or cell-type composition are integrated in the regression model using hidden factors. Those models have been additionally applied to transcriptome analysis [22]. As they do not make specific hypotheses regarding the nature of the data, latent factor models could be applied to any category of association studies regardless of their application field. Method choices and best practices are, however, specific to each field, and have been extensively debated in recent surveys [43, 24].

Most inference methods for latent factor regression models are based on heuristic approaches, lacking theoretical guarantees for identifiability, numerical convergence or statistical efficiency [42]. In addition, existing methods do not always address the confounding problem correctly, building confounder estimates on genetic markers only while ignoring the primary variables. In this study, we propose confounder estimation algorithms that explicitly account for the correlation between confounders and primary variables. The algorithms are based on two distinct regularized least squares methods for *latent factor mixed models*. We present the methods and theoretical developments in the next section. Then we demonstrate that the new methods achieve increased power compared to standard methods in simulations, in an EWAS of patients with rhumatoid arthritis, in a GWAS of patients with celiac disease, and lead to new discoveries in a GEAS of individuals from the 1,000 Genomes Project.

## 2 LFMM algorithms

Consider an *n* × *p* response matrix, **Y**, recording data for *n* individuals. The individual data can correspond to genotypes, methylation profiles or gene expression levels measured from *p* genetic markers or probes. In addition, consider an *n* × *d* matrix, **X**, of individual observations, recording variables of primary interest such as phenotypes or exposure levels. Additional covariates including age and gender of individuals as well as observed confounders could be included in the **X** matrix. Association methods evaluate correlation between the response matrix and the primary variables, and commonly rely on regression models. Latent factor mixed models (LFMMs) are particular regression models defined by a combination of fixed and latent effects [26, 13, 42] as follows

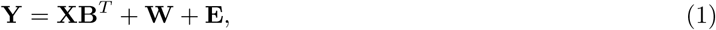

where **W** is an *n* × *p latent matrix* of rank *K*. We defined **U** and **V** as the (unique up to sign) factor and loading matrices obtained from a rank *K* approximation of **W**,

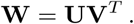

Unobserved confounders are modeled through the *n* × *K* matrix of latent factors, **U**, where the number of confounders, *K*, is determined by model choice procedures (see below). Loadings corresponding to each latent variable are recorded in the **V** matrix, which has dimension *p* × *K*. Fixed effect sizes are recorded in the B matrix, which has dimension *p* × *d*. The **E** matrix represents residual errors, and it has the same dimensions as the response matrix. In this section, we present two statistical learning algorithms for confounder estimation based on *L*^2^ and *L*^1^-regularized least-squares problems.

### *L*^2^-regularized least-squares problem

Statistical estimates of the parameter matrices **U, V, B** in equation

(1) were computed after minimizing the following penalized loss function

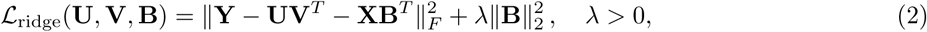

where 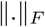 is the Frobenius norm, 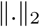 is the *L*^2^ norm, and λ is a regularization parameter. A positive value of the regularization parameter is necessary for identifying the parameter matrices **W** = **UV***^T^* and **B**. To see this, note that for any matrix *P* with dimensions *d* × *p*, we have

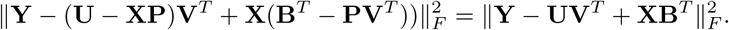

This result entails that the minima of the unregularized (*λ* = 0) least-squares problem are not defined unequivocally, and infinitely many solutions of the least squares problem could exist unless a positive value is considered. As a consequence, any algorithm computing a low rank approximation of a response matrix using their first *K* principal components, and performing a linear regression of the residuals on **X** *d*oes not identify the regression and factor coefficients in equation (1) properly (if we assume Gaussian errors).

### Ridge estimates (LFMM2)

To computer the least-squares estimates of the latent factors, minimization of the 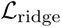 function started with a singular value decomposition (SVD) of the explanatory matrix, **X** = **QΣR***^T^*, where **Q** is an *n* × *n* unitary matrix, R is a *d* × *d* unitary matrix and Σ is an *n* × *d* matrix containing the singular values of **X**, denoted by (*σ_j_*)*_j_*_=1.._*_d_*. The ridge estimates are described as follows

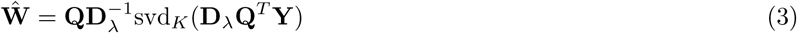

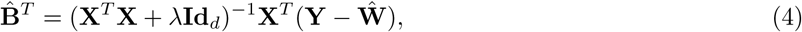

where svd*_K_* (**A**) is the rank *K* singular value decomposition of the matrix **A**, **Id***_d_* is the *d* × *d* identity matrix, and **D**_λ_ is the *n* × *n* diagonal matrix with coefficients defined by

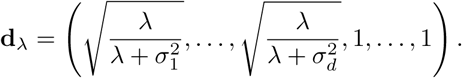

#### Theorem 1.

*Let* λ > 0. *The estimates* **Û**, 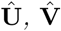 *obtained from the principal component analysis of the matrix* **Ŵ***, and the estimate* 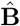 *define a global mimimum of the penalized loss function* 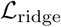.

The proof of Theorem 1 was based on mathematical properties of the SVD, and it can be found in appendix. The result describes a fast algorithm for computing the matrice of confounder estimates, **Û**, with a computing cost determined by the algorithmic complexity of low rank approximation. According to [18], computing **Û** requires *O*(*npK*) operations. This complexity reduces to *O*(*np* log *K*) operations when random projections are used (our implementation). Accounting for the computational cost of *Q^T^Y*, the complexity of the LFMM2 algorithm of order *O*(*n*^2^*p* + *np* log *K*). For studies in which the number of samples, *n*, is much smaller than the number of response variables, *p*, the computing time of ridge estimates is approximately the same as running the low rank approximation SVD algorithm on the response matrix twice.

### *L*^1^-regularized least-squares problem

In addition to the ridge estimates, a sparse regularization approach was considered by introducing penalties based on the *L*^1^ and nuclear norms

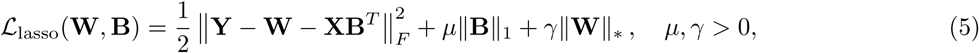

where 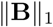 denotes the *L*^1^ norm of **B**, *μ* is an *L*^1^ regularization parameter, **W** is the latent matrix, 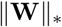 denotes its nuclear norm, and γ is a regularization parameter for the nuclear norm. The *L*^1^ penalty induces sparsity on the fixed effects [40], and corresponds to the prior information that not all response variables may be associated with the primary variables. More specifically, the prior implies that a restricted number of rows of the effect size matrix B are non-zero. The second regularization term is based on the nuclear norm, and it is introduced to penalize large numbers of latent factors. With this term, the 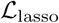 function is a convex function, and convex mimimization algorithms can be applied [31].

### Lasso estimation algorithm (LFMM1)

To simplify the description of the algorithm, let us assume that the explanatory variables, **X**, are scaled so that **X***^T^***X** = **Id***_d_*. Our program implementation is more general (but requires more complex notations). We developed block-coordinate descent methods for minimizing the convex loss function 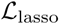 with respect to **B** and **W**. The algorithm is initialized from a null-matrix, **Ŵ**_0_ =0, and iterates the following steps

1. Find 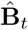 a minimum of the penalized loss function

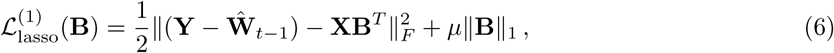

2. Find **Ŵ***_t_* a minimum of the penalized loss function

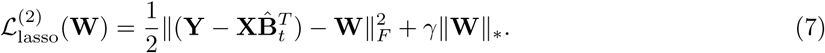

The algorithm cycles through the two steps until a convergence criterion is met or the allocated computing resource is depleted. Each minimization step has a well-defined and unique solution. To see it, note that Step 1 corresponds to an *L*^1^-regularized regression of the residual matrix **Y** – **Ŵ***_t_*_−1_ on the explanatory variables. To compute the regression coefficients, we used Friedman’s block-coordinate descent method [14]. According to [40], we obtained

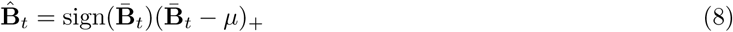

where *s*_+_ = max(0, *s*), sign(*s*) is the sign of *s* and 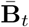 is the classical regression estimate 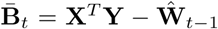. Step 2 consists of finding a low rank approximation of the residual matrix 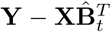 [6]. This approximation starts with a singular value decomposition of the residual matrix 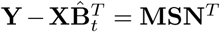, with **M** a unitary matrix of dimension *n* × *n*, **N** a unitary matrix of dimension *p* × *p*, and **S** the matrix of singular values (*s_j_*)*_j_*_=1.._*_n_*. Then, we obtained

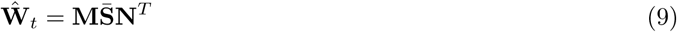

where 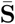 is the diagonal matrix with diagonal terms 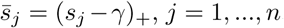. Building on results from [41],the following statement holds.

#### Theorem 2.

*Let μ* > 0 *and γ* > 0. *Then the block-coordinate descent algorithm cycling through Step 1 and Step 2 converges to estimates of* **W** *and* B *defining a global mimimum of the penalized loss function* 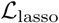.

The proof of Theorem 2 can be found in the appendix. The algorithmic complexities of Step 1 and Step 2 are bounded by a term of order *O*(*pn* + *K*(*p* + *n*)). The computing time of lasso estimates is generally longer than for the ridge estimates, because the LFMM1 algorithm needs to run the SVD and projection steps several times until convergence while the ridge method (LFMM2) requires a single iteration.

### Statistical tests

Suppose we test a single primary variable (*d* = 1, the extension to *d* > 1 variables is straightforward). To test association between **X** and the response variables *Y_j_*, we used the latent score estimates obtained from the LFMM1 or LFMM2 methods as covariates in multiple linear regression models. Our approach is similar to other methods for confounder adjustment in association studies [32, 37, 27, 34, 16]. It differs from other approaches through the latent scores estimates, **Û**, that capture the part of response variation not explained by the primary variable. To test for correlation with the response variable *Y_j_*, we estimated the regression coefficients in a linear regression model

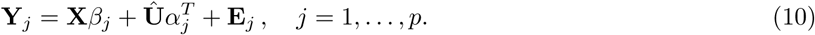

To test the null hypothesis *H*_0_: *β_j_* =0, we used a Student distribution with *n* − *K* − 1 degrees of freedom [19]. To improve test calibration and false discovery rate estimation, we eventually applied an empirical-null testing approach to the test statistics [10].

Remark that in the above equation, causality is modeled when **X** is an exposure variable and **Y** represents a biological measure such as gene expression or DNA methylation levels. When **X** is a phenotypic trait and **Y** represents a biological measure such as a genotype, direct effect sizes can be estimated by switching the response and explanatory variables in the regression model (− = **Y***_j_β_j_*). In addition, tests based on generalized linear models, mixed linear models or robust linear models could be implemented according to similar principles. In the case of mixed linear models, the covariance matrix for random effects can be computed from the *K* estimated factors as *C* = **UU***^T^*/*n*. The methods presented in this study and their extensions were implemented in the R package lfmm.

### R package availability

The R package lfmm is available from the following URL: https://github.com/bcm-uga/lfmm.

## 3 Results and Discussion

### Simulation study

In a series computer experiments, we simulated quantitative trait variables for a worldwide sample of 1,758 human individuals from the 1000 Genomes Project database [1]. Our simulations considered various levels of confounding and numbers of causal variables in the data. For each simulation setting, five data sets were created, representing a total number of 125 data sets. As a baseline, we fitted simple linear regression models (LRM) without adjusting the data for potential confounding effects. This procedure was expected to result in a severe inflation of the test statistic. Six additional association methods were applied to the simulated data: PCA, two variants of SVA, CATE [42] and two variants of LFMM. Additionnally, we implemented an *oracle* method that performed association tests aware of the generating mechanism and confounders. By using the (true) confounders as covariates in its regression model, the oracle method was expected to provide an upper bound on the power of association method that estimates confounding effects (Fig. 1A, Fig. S1). In most simulations, the power of LFMM1, LFMM2 and CATE was identical to the power of the oracle method. PCA (EIGENSTRAT) had genomic inflation factors close to one (Fig. S1), but the power of this method decreased with increasing numbers of causal loci and correlation between confounders and the primary variable. SVA methods had genomic inflation factors close to one for lower levels of confounding (Fig. S1), but their power remained lower than the oracle method for all numbers of causal loci. LFMM1, LFMM2 and CATE acheived substantially more power than SVA1, SVA2 and PCA for higher levels of confounding.

**Figure 1.**
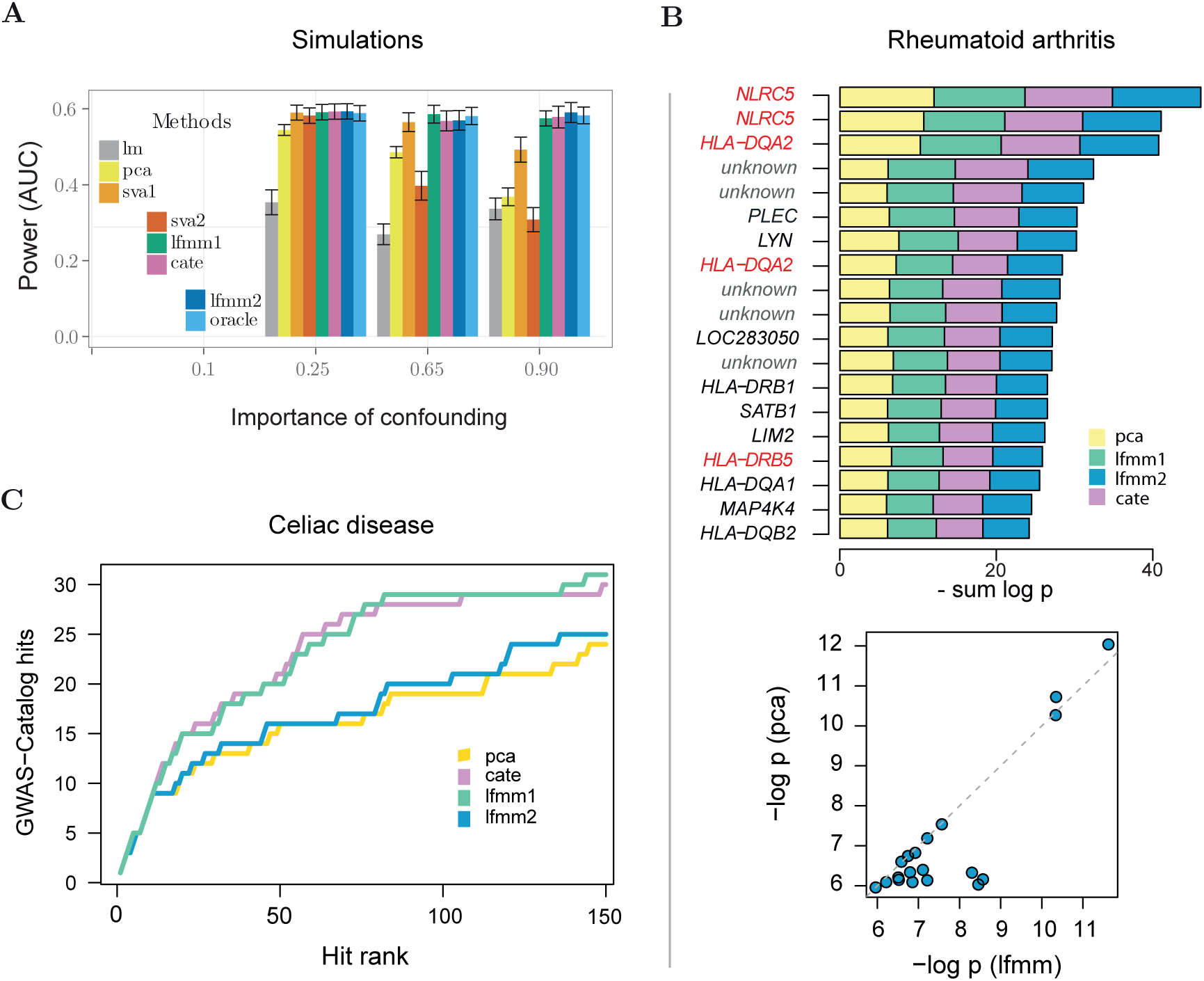
Power of tests. A) Simulations. Power measured by AUC for three levels of confounding. The importance of confounding corresponds to the squared correlation between confounders and primary variables. B) Rheumatoid Arthritis EWAS. Top: List of 19 methylation probes corresponding to the shared top hits of four methods: PCA, CATE, LFMM1 and LFMM2. The list was controlled for a false discovery rate of 1%. Highlighted genes correspond to previously reported discoveries. Bottom: Quantiles plot indicating that PCA correction lead to more conservative tests than methods estimating latent factors. C) Celiac disease GWAS. Ability of four methods to recover genomic regions with known associations with CD. PCA correction led to more conservative tests than methods estimating latent factors.

### Rheumatoid arthritis (RA) EWAS

We performed an EWAS using whole blood methylation data from a study of patients with RA [28]. The cell composition of blood in RA patients is a known source of confounding, and unaccounting for cell type heterogeneity leads to an increased rate of false discoveries [23, 34]. Crossvalidation identified hyperparameter values for the LFMM2 algorithm, and ten confounders were selected (Fig. S2). We implemented five methods for confounder adjustement in EWAS (Fig. 1B, Fig. S3). The resulting discoveries were compared with CpG sites detected by using a reference-based method [20, 45]. PCA, SVA1 and LFMM1 recovered 80% of the reference-based candidates within their eleven top hits, in agreement with the results of a previous analysis with REFACTOR [34]. PCA and SVA1 provided almost identical lists of candidate sites for an expected FDR of 1%. LFMM1 had higher power than PCA and SVA1, and ranked new discoveries above previously discovered candidates (Fig. 1B, Fig. S4). New discoveries included CpG sites in the genes *SPEC* and *LYN* playing an important role in the regulation of innate and adaptive immune responses, and in *HLA-DRB1* having known association with RA [25] (Fig. 1B, Table S1).

### Celiac disease (CD) GWAS

Next we performed a GWAS using SNPs from a study of patients with CD [8]. In GWAS, systematic differences in allele frequencies between patients, known as population structure, are assumed to result in spurious associations and in an increased number of false positive tests [2]. Crossvalidation selected high nine axes of variation in the data (Fig. S5). We implemented four methods for confounder adjustement in GWAS (Fig. 1C, Fig. S5), and we compared the regions found by those methods with the GWAS Catalog for CD. PCA (EIGENSTRAT[32]) led to more conservative tests, resulting in lower power. For LFMM2, the cross-validation method was influenced by SNPs with very large effect sizes in the HLA region, and determined large regularization values and a PCA-like method. Thanks to their robust methods, LFMM1 and CATE had the highest power to detect regions with SNPs included in the GWAS Catalog. Pooling discoveries from all factor methods for an expected FDR level of 1% identified 282 genomic regions, containing 28% of all loci referenced in the GWAS catalog for CD (Fig. 1C, Table 1). For the most powerful method (LFMM1), six genomic regions or loci among the twenty top hits were not referenced in the GWAS catalog [17] (Table 1).

**Table 1.** CD GWAS. Genomic regions corresponding to the first twenty hits of the LFMM1 algorithm (SNP Ids). Rows in bold style correspond to SNPs referenced in the GWAS Catalog for a previously reported association with CD. Chromosome 6 was not included in the analysis.

| Chr | SNP Id | LD block (Mb) | Odds ratio [95% CI] | P value | Q value | Genes |
| --- | --- | --- | --- | --- | --- | --- |
| **3** | **rs1464510** | **189.56-189.61** | **1.30 [1.24-1.36]** | **3.8e-23** | **1.5e-20** | **LPP** |
| **3** | **rs17810546** | **160.99-161.32** | **1.35 [1.26-1.45]** | **1.8e-16** | **6.1e-14** | **IQCJ-SCHIP1, IL12A-AS1, IL12A** |
| **4** | **rs13151961** | **123.19-123.56** | **0.73 [0.68-0.78]** | **1.7e-14** | **5.3e-12** | **KIAA1109, ADAD1** |
| **12** | **rs653178** | **110.25-110.49** | **1.22 [1.16-1.28]** | **6.8e-13** | **2.1e-10** | **CUX2, LINC02356, SH2B3, ATXN2** |
| **2** | **rs917997** | **102.26-102.61** | **1.27 [1.20-1.35]** | **1.5e-12** | **4.6e-10** | **IL1RL1, IL18R1, IL18RAP, MIR4772, SLC9A4, SLC9A2** |
| 4 | rs6840978 | 123.73-123.77 | 0.77 [0.72-0.82] | 1.2e-11 | 3.5e-09 | IL21-AS1 |
| 3 | rs9811792 | 161.12-161.18 | 1.18 [1.12-1.24] | 6.6e-11 | 1.9e-08 | IL12A-AS1 |
| **3** | **rs13098911** | **45.98-46.21** | **1.32 [1.22-1.43]** | **2.1e-10** | **5.8e-08** | **FYCO1, FLT1P1, CCR3** |
| **1** | **rs2816316** | **190.77-190.80** | **0.78 [0.72-0.83]** | **2.2e-10** | **6.2e-08** |  |
| **3** | **rs6441961** | **46.26-46.33** | **1.21 [1.15-1.27]** | **1.7e-08** | **4.6e-06** | **CCR3, UQCRC2P1** |
| **2** | **rs4675374** | **204.29-204.52** | **1.23 [1.16-1.31]** | **2.1e-07** | **5.4e-05** | **CD28, ICOS** |
| 2 | rs1018326 | 181.54-181.78 | 1.15 [1.10-1.21] | 4.4e-07 | 1.1e-04 | UBE2E3, LINC01934 |
| 3 | rs7648827 | 46.56-46.56 | 1.22 [1.12-1.33] | 4.6e-07 | 1.2e-04 | LRRC2 |
| **2** | **rs13003464** | **60.95-61.24** | **1.19 [1.13-1.25]** | **5.0e-07** | **1.3e-04** | **LINC01185, REL, PUS10, RNA5SP95, KIAA1841, C2orf74** |
| **10** | **rs1250552** | **80.71-80.74** | **0.84 [0.80-0.88]** | **5.2e-07** | **1.3e-04** | **ZMIZ1** |
| 3 | rs7629708 | 189.56-189.62 | 1.17 [1.11-1.24] | 1.0e-06 | 2.5e-04 | LPP |
| **22** | **rs2298428** | **20.13-20.31** | **1.17 [1.10-1.24]** | **1.3e-06** | **3.3e-04** | **HIC2, UBE2L3, YDJC, CCDC116** |
| 18 | rs1394466 | 48.93-49.30 | 1.14 [1.08-1.20] | 1.5e-06 | 3.6e-04 | DCC |
| **18** | **rs1893217** | **12.80-12.84** | **1.16 [1.08-1.23]** | **1.6e-06** | **4.0e-04** | **PTPN2** |
| **1** | **rs864537** | **165.66-165.70** | **0.87 [0.83-0.92]** | **1.7e-06** | **4.2e-04** | **POU2F1, CD247** |

### Human GEAS

To detect genomic signatures of adaptation to climate in humans, we performed a GEAS using 5,397,214 SNPs for 1,409 individus from the 1,000 Genomes Project [1], and bioclimatic data from the WorldClim database [12] (Fig. S6). Nine confounders were estimated by LFMM2, mainly describing correlation between population structure and climate in the sample (Fig. 2A, Fig. S7, Fig. S8). Four methods for confounder adjustement led to a list of 836 (1335) SNPs after pooling the list of candidates from the four methods (expected FDR = 1%-5%). A variant prediction analysis reported a large number of SNPs in intergenic and intronic regions, with an over-representation of genic regions (Fig. 2B). Top hits represented genomic regions important for adaptation of humans to environmental conditions. The hits included functional variants in the *LCT* gene, and SNPs in the *EPAS1* and *OCA2* genes previously reported for their role in adaptation to diet, altitude or in eye color [11] (Fig. 2B, Fig. S9, Table S2).

**Figure 2.**
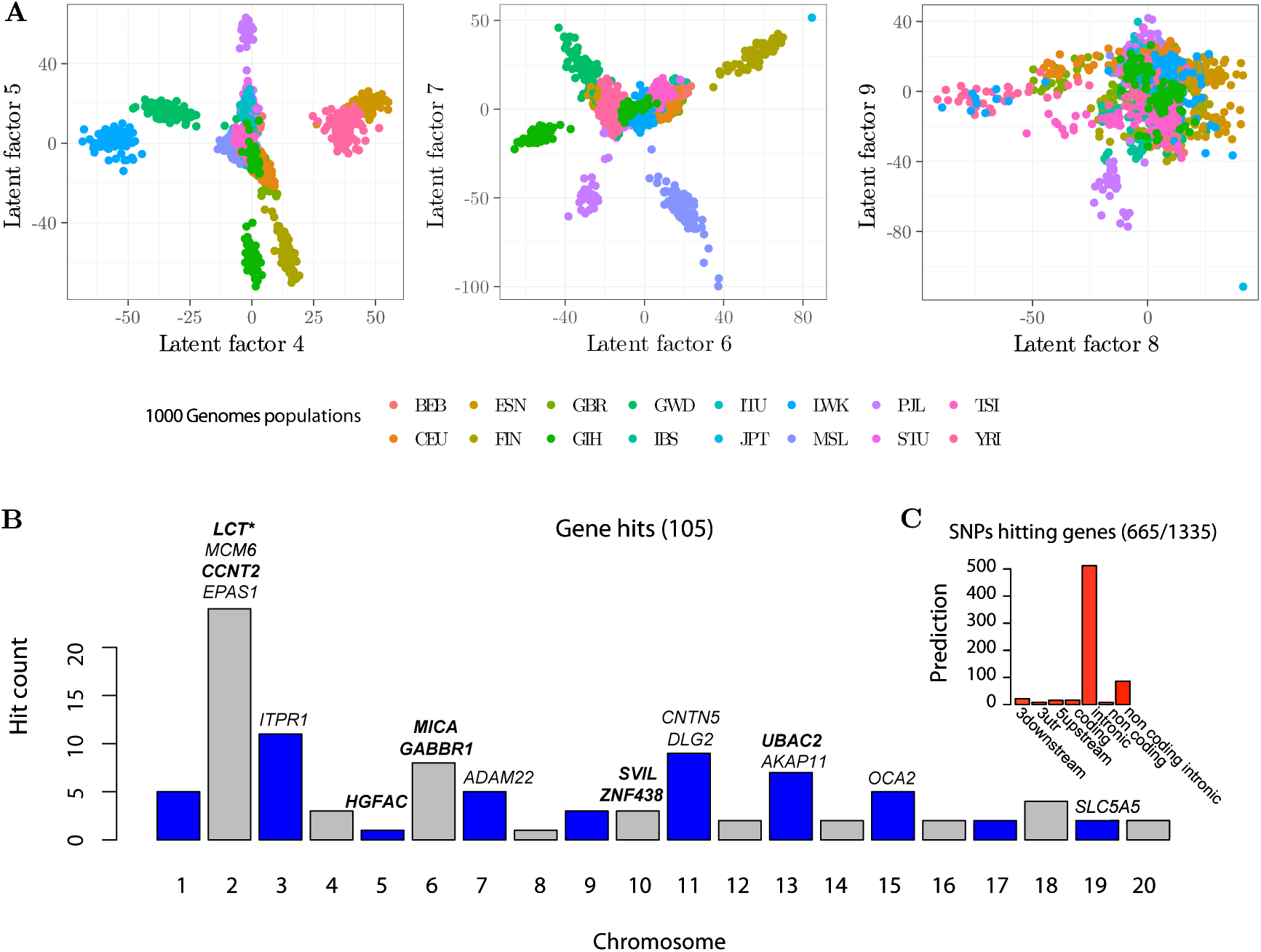
Gene environment association study. Association study based on genomic data from the 1000 Genomes Project database and climatic data from the Worlclim database. A) Latent factors estimated by LFMM2. B) Target genes corresponding to top hits of the GEAS analysis (expected FDR level of 5%). The highlighted genes correspond to functional variants. Predictions were obtained from the variant effect predictor program.

### Conclusions

In this study we introduced two statistical learning algorithms for confounder estimation based on *L*^2^ and *L*^1^-regularized least-squares problems for latent factor regression models. We used those algorithms for testing associations between a response matrix **Y** and a primary variable matrix **X** in EWAS, GWAS and GEAS. In those applications, standard association methods have mainly focused on corrections for specific confounding effects such as individual relatedness or cell type composition. In contrast, LFMMs do not put prior information on any particular source of confounding, but account for correlation between confounders and primary variables. Compared to PCA approaches, LFMMs gained power by removing the part of genetic variation that could not be explained by the primary variables [13]. In GWAS, LFMM extends tests performed by the EIGENSTRAT program by improving estimates of principal components [32]. For EWAS, LFMMs extended surrogate variable analysis (SVA) [26] also achieving an increased power. In comparison with other algorithms, LFMM1 and LFMM2 have mathematical guarantees to provide globally optimal solutions of least-squares estimation problems.

Like several factor methods, the computational speed of LFMM methods is mainly influenced by the algorithmic complexity of low rank approximation of large matrices. The algorithmic complexity of LFMM methods in similar to PCA or SVA, of order *O*(*np* log *K*) for LFMM1 and *O*(*n*^2^*p* + *np* log *K*) for LFMM2. LFMM2 is generally faster than LFMM1 because all computations are based on a unique round of SVDs. These approaches are faster than algorithms based on mixed linear models [44] and faster than Bayesian methods currently used in GEAS [13]. Although potential improvements such as random effects, logistic or robust regressions and stepwise conditional tests were not included in our results, those options are available with the lfmm program, and they may provide additional power to detect true associations.

## 4 Materials and Methods

### Multiple linear regression with principal components

We implemented a standard approach that estimates confounders from the PCA of the response matrix **Y**. Scaling the response matrix, this approach is similar to the EIGENSTRAT method [32]. As a baseline, we also implemented linear regression models (LRM) that did not include correction for confounding. In the presence of confounding, LRM are expected to lead to inflation of false positive tests, whereas the PCA approach is expected to lead to overly conservative tests.

### Surrogate variable analysis

Surrogate variable analysis (SVA) was introduced to overcome the problems caused by heterogeneity in gene expression studies. SVA is based on the latent factor regression model presented in this study, and is potentially useful for confounder adjustment in any type of genome-wide association studies. Two distinct SVA algorithms were implemented SVA1 [26] and SVA2 [27]. In a first step, the SVA1 algorithm estimates loadings of a PCA of the residuals of the regression of the response matrix **Y** on **X**. The second step of the SVA1 algorithm determines a subset of response variables exhibiting low correlation with **X**, and uses this subset of variables to estimate the score matrix. SVA1 is similar to LFMM2 with regularization term set to λ = 0, a degenerate case for the least-squares problem. The SVA2 method is an iterative approach. In SVA2, the second step of SVA1 is modified so that a weight is given to each response variable. Weights are used to compute a weighted PCA of the regression residuals, and the cycle is iterated until a convergence criterion is met. The SVA methods were implemented by using the R package sva [27].

### Confounder adjusted testing and estimation

Confounder adjusted testing and estimation (CATE) [42] is a recent estimation method based on latent factor regression models. CATE uses a linear transformation of the response matrix such that the first axis of this transformation is colinear to **X**. CATE and LFMM2 apply different transformations to the response matrix, but the CATE estimates are comparable to LFMM2 estimates (although CATE estimates do not solve a least-squares problem). Asymptotic results obtained for CATE estimates were also valid for LFMM2 estimates. The CATE method was implemented in the R package cate [42].

### Simulated data

We used empirical data from a world-wide sample of 1,758 human genotypes from the 1,000 Genomes Project [1] for simulating quantitative phenotypes, **X**, latent factors **U** and a response matrix **Y**. For the simulation, an initial data matrix, **Y**^⋆^, including 52,211 single nucleotide polymorphisms (SNPs) from chromosomes 1 and 2 was considered. To create *K* articifial confounders, we first performed a PCA of the **Y**^⋆^ matrix, and retained *K* = 5 principal components. The eigenvalues, 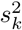, were computed for each retained component. Then a primary variable **X** and five latent variables latentes, **U**, were simulated by using a multivariate Gaussian distribution

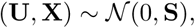

where **S** was the covariance matrix defined by

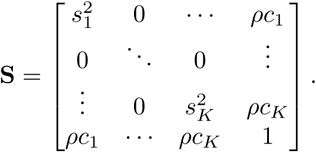

The *c_k_* coefficients were sampled from a uniform distribution taking values in the range (−1, 1), and *ρ* was inversely proportional to the square root of 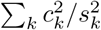 (which was less than one). The coefficient of proportionality was chosen so that the percentage of variance of **X** explained by the latent factors ranged between (0.1, 1). The effect size matrix, **B**, was generated by setting a proportion of effect sizes to zero. Non-zero effect sizes were sampled according to a standard Gaussian distribution *N*(0, 1). The proportion of null effect sizes ranged between 80% and 99%. We eventually created a response matrix, **Y**, by simulating from the generative model of the latent factor model as follows

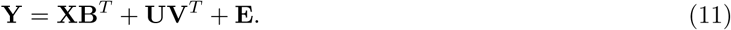

In those simulations, *K* latent variables had the same variance as original PCs from the 1,000 Genomes Project data set, and we controlled the correlation between simulated phenotypes (primary variables) and confounders.

To evaluate the capabilities of methods to identify true positives, we used the area under the precision-power curve (AUC) as a global estimate of power. Precision is the proportion of true positives in a candidate list of positive tests. Power is the number of true positives divided by the number of true associations. To evaluate whether the methods have inflated number of false positives, we computed a genomic inflation factor using the median of squared z-scores divided by the median of the chi-squared distribution with one degree of freedom [5].

### Rheumatoid arthritis (RA) data set

We performed an EWAS using whole blood methylation data from a study of patients with rheumatoid arthritis [28, 34, 45]. The RA data are publicly available and were downloaded from the GEO database (accession number GSE42861). For this study, beta-normalized methylation levels at 485,577 probed CpG sites were measured for 354 cases and 335 controls [28]. Following [45], probed CpG sites having a methylation level lower than 0.2 or greater than 0.8 were filtered out. Then, the data were centered and scaled for a standard deviation of one. Since the cell composition of blood in RA patients typically differs from that in the general population, there is a risk for false discoveries that stem from unaccounted-for cell type heterogeneity [34]. Age, gender and covariates such as tobacco consumption may also have significant effects on DNA methylation. To evaluate whether the methods presented here can correct confounding due to those factors, we did not include them as covariates in regression analyses. Seven EWAS methods were applied to the RA data set, including LRM, PCA, two variants of SVA, CATE and two variants of LFMM. Candidate lists of CpG sites were controlled for a false discovey rate of 1% after recalibration of the test significance values, and compared to the candidates obtained with a reference-based method and controlling for age, gender and smocking status. FDR control was implemented through the qvalue function of the R program [38].

### Celiac disease (CD) data set

We performed a GWAS using SNPs from a study of patients with celiac disease [8]. The CD data were downloaded from the Wellcome Trust Case Control Consortium https://www.wtccc.org.uk/. For this study, SNP genotypes were recorded at 485,577 loci for 4,496 cases and 10,659 controls. The genotype matrix was preprocessed so that SNPs with minor allele frequency lower than 5% and individuals with relatedness greater than 8% were removed from the matrix. We used the program BEAGLE to impute missing data in the genotype matrix [4]. We performed LD pruning to retain SNPs with the highest frequency in windows of one hundred SNPs. The filtering steps were implemented in the PLINK software [33], and resulted in a subset of 80,275 SNPs. Five GWAS methods were applied to the CD data set: LRM, PCA (EIGENSTRAT), CATE, and two LFMM estimation algorithms. For the last four methods, the confounders were identified based on the 80,275 pruned genotypes. The tests were performed on the full set of imputed genotypes, and the SNP positions were grouped into clumps of correlated SNPs, using the clumping algorithm implemented in PLINK. The significance value for a clump of SNPs was considered to be the lowest value among all positions. FDR control was applied on the clumped significance values using qvalue. Candidates resulting from the five analyses were compared to the GWAS catalog for known association with CD [29]. Chromosome 6, which contained the strongest association signals with CD, was treated separately. For this chromosome, all methods performed equally well at detecting six SNPs from the HLA locus referenced in the GWAS catalog.

### Gene-environment association study

We performed a GEAS using whole genome sequencing data and bioclimatic variables to detect genomic signatures of adaptation to climate in humans. The data are publicly available, and they were downloaded from the 1,000 Genomes Project phase 3 [1] and from the WorldClim database [12]. The genomic data included 84.4 millions of genetic variants genotyped for 2,506 individuals from 26 world-wide human populations. Nineteen bioclimatic data were downloaded for each individual geographic location, considering capital cities of their country of origin. The bioclimatic data were summarized by projection on their first principal component axis. The genotype matrix was preprocessed so that SNPs with minor allele frequency lower than 5% and individuals with relatedness greater than 8% were removed from the matrix. Admixed individuals from Afro-american and Afro-Caribbean populations were also removed from the data set. After those filtering steps, the response matrix contained 1,409 individus and 5,397,214 SNPs. We performed LD pruning to retain SNPs with the highest frequency in windows of one hundred SNPs, and identified a subset of 296,948 informative SNPs. Four GEAS methods were applied to the 1,000 Genomes Project data set: PCA (EIGENSTRAT), CATE, and two LFMM estimation algorithms. For all methods the latent factors were estimated from the pruned genotypes, and association tests were performed for all 5,397,214 loci. Candidates obtained from clumps with an expected FDR level of 1% were analyzed using the Variant Effect Predictor (VEP) program [30].

### Cross-validation and model choice

Choosing regularization parameters and the number of latent factors can be achieved by using cross-validation methods. We developed a cross-validation approach appropriate to latent factor regression models. Cross-validation partitions the data into a training set and a test set. The training set is used to fit model parameters, and prediction errors can be measured on the test set. In our approach, the reponse and explanatory variables were partitioned according to their rows (individuals). We denote by *I* the subset of individual labels on which prediction errors are computed. Estimates of effect sizes, 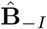, and factor loadings, 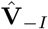, were obtained from the training set. Next, the set of columns of the response matrix were partitioned. Denoting by *J* the subset of columns on which the prediction errors were computed, a score matrix was estimated from the complementary subset as follows

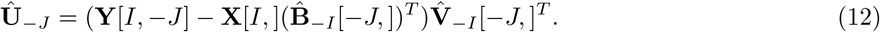

In this notation, the brackets indicate which subsets of rows and columns of a matrix were selected. A prediction error was computed as follows

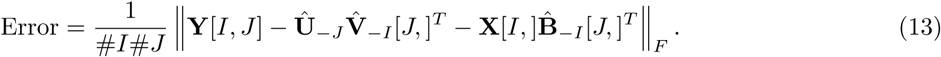

Parameters leading to the lowest prediction errors were retained for data analysis.

Additional heuristics were used to determine the number of latent factors and the nuclear norm parameter for LFMM1 latent matrix estimates. For choosing the number of latent factors, *K*, we considered the matrix **D**_λ_ defined in the statement of Theorem 1, and the **Q** unitary matrix obtained from the SVD of **X**. The number of latent factors, *K*, was estimated after a spectral analysis of the matrix **D**_0_**Q***^T^***Y**. We determined it by estimating the number of components in a PCA of the matrix **D**_0_**Q***^T^***Y**. In our experiments, we used the “elbow” method based on the scree-plot of PC eigenvalues. Estimated values for *K* were confirmed based on prediction errors computed by cross-validation. The *L*^1^-regularization parameter, *μ*, was determined after the proportion of non-zero effect sizes in the **B** matrix, which was estimated by cross-validation. Having set the proportion of non-zero effect sizes, *μ* was computed by using the regularization path approach proposed in [15]. The regularization path algorithm was initialized with the smallest values of *μ* such that

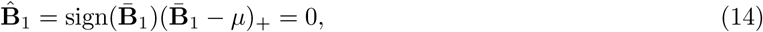

where 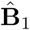 resulted from Step 1 in the lasso (LFMM1) estimation algorithm. Then, we built a sequence of *μ* values that decreases from the inferred value of the parameter, *μ*^max^, to *μ*^min^ = ∊*μ*^max^. We eventually computed the number of non-zero elements in 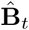, and stopped when the target proportion was reached. The nuclear norm parameter (*γ*) determines the rank of the latent matrix **W**. We used a heuristic approach to evaluate γ from the number of latent factors *K*. The singular values (*λ*_1_&*,λ_n_*) of the response matrix **Y** were computed, and we set

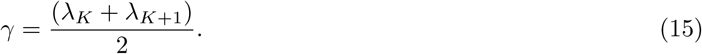

In our experiments, the lasso estimation algorithm always converged to a latent matrix estimate having rank *K*.

## Appendix

In this section, we provide proofs for Theorems 1 and 2. For Theorem 1, we define the singular value decomposition (SVD) of the explanatory matrix, as **X** = **QΣR***^T^*, where **Q** is an *n* × *n* unitary matrix, **R** is a *d* × *d* unitary matrix and **Σ** is an *n* × *d* matrix containing the singular values of **X**, (*σ_j_*)*_j_*_=1_..*_d_*. Let *λ* > 0, and the estimates given by

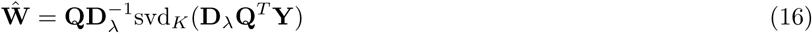

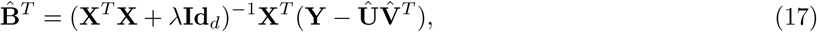

where **D***_λ_* is the *n* × *n* diagonal matrix with diagonal terms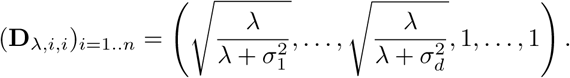

We prove that those estimates define a global mimimum of the function 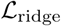.

### Proof.

If we assume **U** are **V** to be known, then the 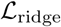 function is convex with respect to the variable **B**. A global minimum for this variable can be found by computing the derivative of 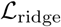 with respect to **B** and setting it to zero. This leads to the following solution

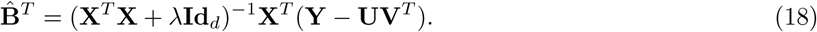

This solution is merely the ridge estimate for a linear regression of the response matrix **Y** − **UV***^T^* on **X**. Thus, the problem amounts to minimizing the (implicit) function

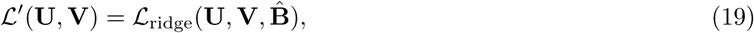

where 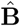 was defined above. Consider the SVD of **X**

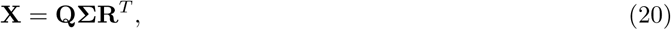

where **Q** is a unitary matrix of dimensions *n* × *n*, **R** is a unitary matrix of dimensions *d* × *d*, and **Σ** is a matrix of dimensions *n* × *d* containing the singular values (*σ_j_*)*_j_*_=1_..*_d_*. The *L* function rewrites as follows

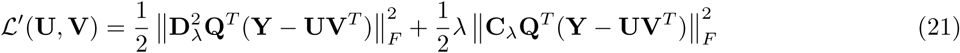

where C_A_ is a matrix of dimensions d × *n*, with zero coefficients except for the diagonal terms

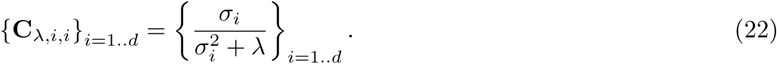

The D_a_ is a diagonal matrix of dimensions *n* × *n* such that

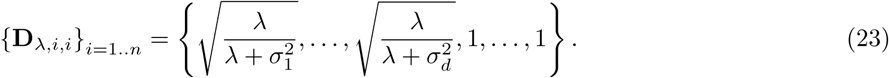

Direct calculus shows that we have

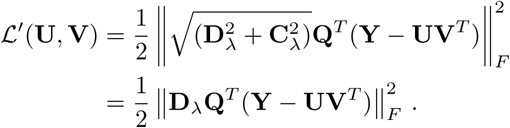

This equation shows that mimimizing the objective function 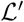 is equivalent to finding the best approximation of rank *K* for **D***_λ_***Q***^T^***Y**. According to [9], this solution is given by the rank *K* SVD of **D***_λ_***Q***^T^***Y**. Eventually, this concludes the proof that

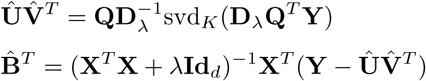

defined a global minimum for the 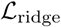 function. □

Now, turn to the proof of Theorem 2. Let *μ* > 0 and *γ* > 0. Then Theorem2 states that the block-coordinate descent algorithm converges to estimates of **W** and **B** defining a global mimimum of the function 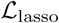.

### Proof.

The result is a consequence of the convexity of 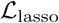 function, and the fact that we can write

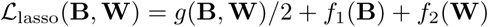

where 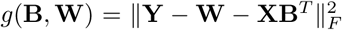 is a differentiable convex function, and 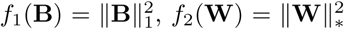 are continuous convex functions. The proof of Theorem 2 relies on the following proposition adapted from [3] and [41].

#### Proposition.

Let *A* = *A*_1_ × *A*_2_ × … × *A_m_* be a Cartesian product of closed convex sets. Consider a continuous convex function *f* defined on *A* as follows

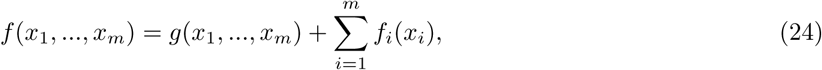

where g is a differentiable convex function, and for all *i*, *f_i_* is a continuous convex function. Let (*x^t^*^+1^) be the sequence of values defined by the block-coordinate descent algorithm

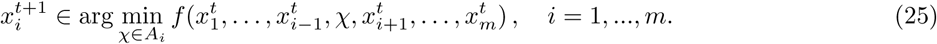

Then a limit point of (*x^t^*) defines a global minimum of *f*. □

## Accession codes

SNP genotypes used in our simulation analysis are publicly available and were downloaded from the 1000 Genome Project database. The RA data are publicly available and were downloaded from the GEO database (accession number GSE42861). The CD data are publicly available and were downloaded from the Welcome Trust Case Control Consortium database (agreement number 1248).

## Acknowledgments

This work has been supported by a grant from LabEx PERSYVAL Lab, ANR-11-LABX-0025-01, funded by the French program Investissement d’Avenir.

## Author contributions

K.C. performed research, contributed analytic tools, and analyzed data. O.F. designed research, contributed analytic tools, analyzed data, and wrote the paper.

